# Quantized multi-task learning for context-specific representations of gene network dynamics

**DOI:** 10.1101/2024.08.16.608180

**Authors:** Han Chen, Madhavan S. Venkatesh, Javier Gómez Ortega, Siddharth V. Mahesh, Tarak N. Nandi, Ravi K. Madduri, Karin Pelka, Christina V. Theodoris

## Abstract

While often represented as static entities, gene networks are highly context-dependent. Here, we developed a multi-task learning strategy to yield context-specific representations of gene network dynamics. We assembled a corpus comprising ∼103 million human single-cell transcriptomes from a broad range of tissues and diseases and performed a two stage pretraining, first with non-malignant cells to generate a foundational model and then with continual learning on cancer cells to tune the model to the cancer domain. We performed multi-task learning with the foundational model to learn context-specific representations of a broad range of cell types, tissues, developmental stages, and diseases. We then leveraged the cancer-tuned model to jointly learn cell states and predict tumor-restricting factors within the colorectal tumor microenvironment. Model quantization allowed resource-efficient fine-tuning and inference while preserving biological knowledge. Overall, multi-task learning enables context-specific disease modeling that can yield contextual predictions of candidate therapeutic targets for human disease.

---

Mapping gene regulatory networks in development and disease enables the discovery of key network regulators and network-correcting therapies that restore disease-dependent networks back to the normal state^1,2^. However, mapping the gene network architecture using traditional methods requires large amounts of transcriptomic data to learn the connections between genes, impeding discoveries in settings with limited data, including rare diseases and diseases affecting clinically inaccessible tissues. Yet, advances in sequencing technologies have driven a rapid expansion in the amount of single-cell transcriptomic data available from tissues more broadly. Standard approaches using task-specific data to train a computational model to make predictions in that particular task require retraining from scratch with new task-specific data for each new task, therefore not fully taking advantage of this broader available data. On the other hand, the machine learning approach of transfer learning leverages large-scale general datasets to pretrain models to gain foundational knowledge that can then be transferred to a vast array of downstream tasks, enabling predictions with little or no task-specific training data^3-5^.

We previously developed a transfer learning strategy for network biology, pretraining a foundational deep learning model, Geneformer, on ∼30 million single-cell transcriptomes to gain a fundamental understanding of network dynamics^6^. We demonstrated that this approach was able to drive biological insights that were experimentally verified with functional assays in cells. For example, Geneformer discovered a novel transcription factor in cardiomyocytes with zero-shot learning and predicted candidate therapeutic targets for cardiomyopathy that improved contractility in an induced pluripotent stem cell (iPSC) model of the disease^6^.

Overall, there has been a recent growth in the adoption of transfer learning for network biology, and multiple foundation models have been pretrained using large-scale single-cell -omics data to enable predictions in a diverse array of downstream tasks^6-12^. Many of these foundation models, like Geneformer, employ a transformer^3^ architecture, which yields context-aware embeddings and predictions. Context-aware approaches are critical for modeling gene regulatory networks, which are highly dependent on cell type, tissue, disease state, and developmental and aging contexts. However, standard single-task fine-tuning, such as learning cell types within normal tissues, or learning disease states within a single cell type, limits the biological dimensions from which the model can learn in a unified manner and may unintentionally collapse biologically meaningful variation within the embedding space.

Here, we developed a multi-task learning strategy to yield context-specific representations of gene network dynamics across cell types, tissues, diseases, and developmental stages. We assembled a large-scale pretraining corpus, Genecorpus-103M, comprising ∼103 million human single-cell transcriptomes from a diverse range of tissues and disease states from publicly available data. We performed an initial self-supervised pretraining with ∼95 million cells excluding cells with high mutational burdens (e.g. malignant cells and immortalized cell lines) and using an expanded input size of 4096 to model a larger context of genes per cell. Pretraining with the larger, more diverse corpus, increased model parameters, and expanded input size boosted zero-shot predictions in a diverse set of downstream tasks. We then performed multi-task fine-tuning to jointly learn cell types, tissues, disease states, and developmental stages, yielding context-specific representations of gene network dynamics across these biological dimensions. Because the initial pretraining excluded malignant cells, we designed a strategy for domain-specific continual learning to tune the model with ∼14 million cells from a broad range of cancer studies. We then leveraged this cancer-tuned model to jointly learn cell states within the tumor microenvironment and predict factors that would shift cells to a tumor-restricting or immune-activating state using in silico treatment analysis. Furthermore, we demonstrated that model quantization allows resource-efficient finetuning and inference while preserving biological knowledge. Overall, multi-task learning represents an effective method for jointly learning multiple biologically informative features to yield context-specific representations of gene network dynamics and predict context-specific therapeutic targets for diseases with multicellular pathology.

## Results

### Pretraining with larger and more diverse corpus enabled predictions in previously elusive tasks

We previously reported that increasing the size and diversity of the pretraining corpus for Geneformer consistently improved the model’s predictive potential^6^. Since the pretraining of Geneformer in June 2021, there has been a significant expansion in both the amount and diversity of publicly available human single-cell transcriptomic data, suggesting we could use this data to now train an even more effective foundational model. Therefore, we expanded our corpus to ∼103 million human single-cell transcriptomes from an even more diverse array of tissue and disease contexts (Fig. 1a-b, Extended Data Table 1, Extended Data Fig. 1a-c). We balanced the data such that no tissue composed more than 25% of the data and performed scalable quality control filtering. We also performed deduplication of studies by DOI to preclude training with duplicated cells, which can significantly overestimate corpus size due to studies being deposited in multiple databases (Extended Data Fig. 1a). Because the technology has advanced since 2021 with more genes now being detected per cell that is sequenced, we expanded the model’s input size to 4096, which fully encompasses 93% of the cells in the pretraining corpus (Extended Data Fig. 1d). Due to the quadratic time complexity of dense attention, this doubling of the input size increased the computational intensity quadratically, but allowed the model to learn from a larger gene network context for each cell.

**Fig. 1.**
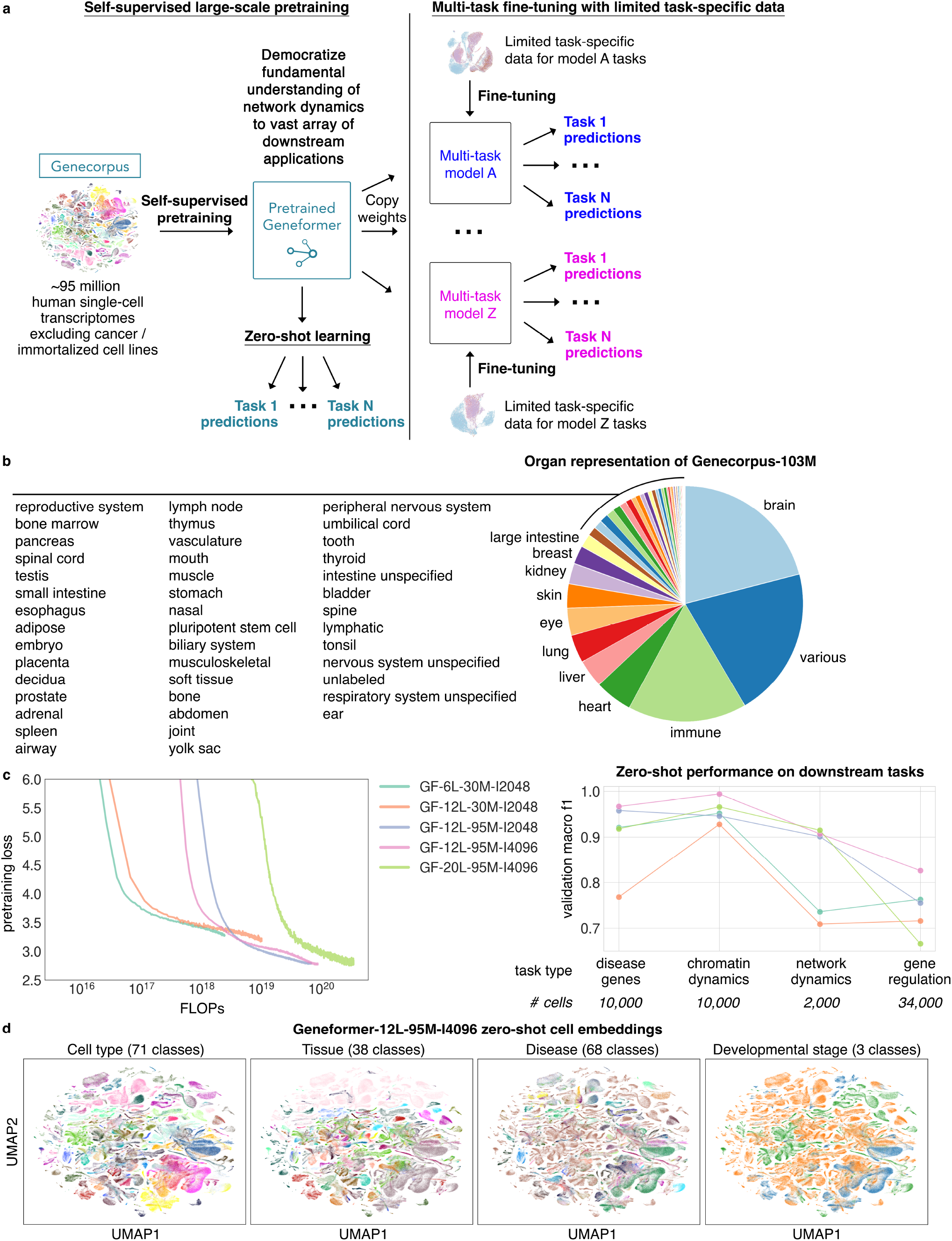
Geneformer transfer learning strategy. **a**, Initial self-supervised, large-scale pretraining on a generalizable learning objective yields a pretrained model with a fundamental understanding of network dynamics. This baseline knowledge can be democratized to a vast array of downstream applications either through zero-shot learning, where the pretrained model is used directly without fine-tuning, or with fine-tuning, where the model learns from limited task-specific data to make much better predictions in the downstream tasks compared to if the model used that limited data alone without the fundamental knowledge gained during the large-scale pretraining. Multi-task fine-tuning enables the model to learn context-specific representations of cell states from multiple cross-informative biologically-relevant tasks. **b**, Organ representation of Genecorpus-103M. **c**, Pretraining loss (*left*) and zero-shot performance on a diverse panel of downstream tasks (*right*) for each pretrained model (GF=Geneformer, L=Layers, M=Million cells, I=Input size). Number of cells for the downstream tasks indicates the number of cells from which zero-shot gene embeddings were extracted for classification. **d**, Zero-shot cell embeddings from GF-12L-95M-I4096 for 779,905 representative cells from the CELLxGENE corpus balanced across cell types, tissues, diseases, and developmental stages and colored by consolidated labels for those cell attributes.

We then pretrained an updated Geneformer model using the larger input size of 4096 per cell and the expanded pretraining corpus with ∼128 billion tokens, where each gene is a token within the dictionary of 20,271 genes. Each cell was presented to the model as a rank value encoding, as previously described^6^. For this primary pretraining stage we used ∼95 million human transcriptomes, excluding cells with high mutational burdens such as malignant cells and immortalized cell lines. We excluded these cells that may have a high abundance of gain of function mutations that may lead to genes having an unpredictably different function than what the model would interpret from other cells with low mutational burdens when observing only transcriptomic data without accompanying genomic sequencing. To match the increase in pretraining data, we also increased the depth of the model, maintaining the width-to-depth aspect ratio, and compared the pretraining loss per computation and tokens observed by the model. Pretraining loss improved with increasing floating point operations (FLOPs), though the largest 20 layer model did not surpass the intermediate-sized 12 layer model until nearly three epochs of training (Fig. 1c, Extended Data Fig. 1e).

The updated models demonstrated enhanced zero-shot learning capabilities across a diverse panel of biologically meaningful downstream tasks in the domains of disease genes, chromatin dynamics, network dynamics, and gene regulation (Fig. 1c). The updated model enabled predictions in previously elusive tasks, such as understanding whether transcription factors act in short or long range with their targets, which is especially difficult to ascertain using only transcriptomic data as input with no information about genomic distance. At the cell level, the zero-shot embeddings captured multiple dimensions of biologically meaningful attributes, including cell type, tissue origin, developmental stage, and disease status (Fig. 1d). Overall, increasing the size and diversity of the pretraining data as well as the model size significantly improved performance in a diverse panel of biologically meaningful downstream tasks.

### Model quantization allowed resource-efficient fine– tuning with nearly equivalent predictive accuracy

Although fine-tuning is generally much less computationally intensive than pretraining, hyperparameter tuning (optimizing the settings that control the model learning process) can rapidly increase the resource requirements due to the need for repeated training attempts to search the space of possible settings. This can impede the accessibility of the model in settings with low GPU resources. To address this, we tested model quantization to 4-bit precision using Quantized Low Rank Adapters (QLoRA)^13^. This approach backpropagates gradients through the frozen, 4-bit quantized Geneformer into low rank adapters to reduce memory usage and training time.

To test the benefit of quantization, we used the largest 20 layer model that requires more compute at baseline. Fine-tuning the 4-bit quantized Geneformer with a relatively low adapter rank of 16 was sufficient to preserve full 32-bit fine-tuning performance in tasks across the four biologically diverse domains while reducing memory by 1/3 and taking 1/3 the time to train with the same batch size as the 32-bit model (Fig. 2). Of note, because the memory usage is lower, the true maximal time-savings are significantly larger since larger batches of data can be run through the 4-bit model at the same memory scale.

**Fig. 2.**
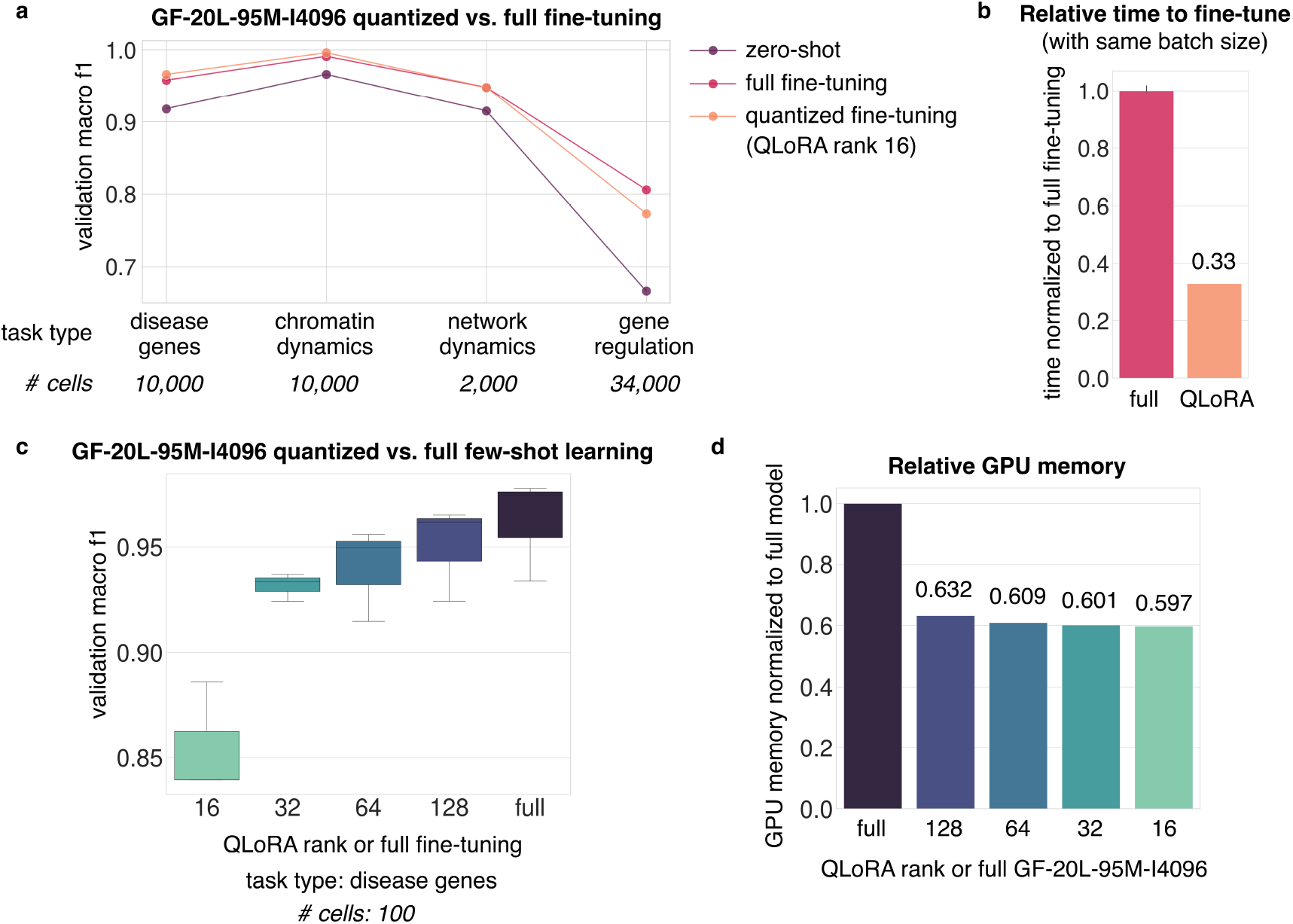
Model quantization enabled resource-efficient fine-tuning. **a**, Validation macro F1 on a diverse panel of downstream tasks for GF-20L-95M-I4096 by zero-shot learning or full fine-tuning or fine-tuning of the quantized 20 layer model with QLoRA rank 16. Number of cells for the downstream tasks indicates the number of cells from which zero-shot gene embeddings were extracted for classification or number of cells used as examples for fine-tuning. **b**, Relative time to fine-tune for full fine-tuning of GF-20L-95M-I4096 vs. fine-tuning the quantized 20 layer model. Of note, time was quantified per the same batch size, but quantized fine-tuning would take even less time in actuality because the lower memory requirements of the model would allow larger batch sizes. **c**, Validation macro F1 on disease genes downstream task (distinguishing dosage-sensitive vs. -insensitive transcription factors) with few-shot learning with only 100 example cells for GF-20L-95M-I4096 by full fine-tuning or fine-tuning the quantized 20 layer model with increasing QLoRA rank. **d**, Relative GPU memory usage of GF-20L-95M-I4096 vs. the quantized 20 layer model with varying QLoRA ranks.

We then tested the ability of the 4-bit quantized model to replicate full fine-tuning in the few-shot setting (Fig. 2c-d). Fine-tuning the 32-bit 20 layer model with just 100 cells was sufficient to yield accurate predictions of dosage-sensitive transcription factors, demonstrating the ability of the model to learn from increasingly limited task-specific data. However, in the few-shot setting, the relatively low adapter rank of 16 was not able to match the performance of full fine-tuning, and increasing the adapter rank was necessary to closely approach the performance of full fine-tuning. Nevertheless, the 128-rank quantized model continued to provide roughly equivalent time and memory savings as the 16-rank version, with no change in time requirements and very minimal change in memory usage. Overall, quantization of Geneformer enabled resource-efficient fine-tuning with nearly equivalent predictive accuracy.

### Multi-task learning strategy yielded context-specific representations of cell states

Fine-tuning provides a valuable method to instruct the model about a specific gene characteristic or cell state, such as distinguishing dosage-sensitive genes or disease states. For example, we previously fine-tuned Geneformer-6L-30M to distinguish cardiomyocytes from non-failing hearts vs. hearts affected by dilated or hypertrophic cardiomyopathy. We then leveraged this finetuned model to perform in silico treatment analysis that predicted therapeutic targets which we experimentally validated to improve the contractility of cardiac microtissues in an induced pluripotent stem cell (iPSC) model of the disease^6^.

However, many diseases are affected by multicellular pathologies where jointly learning about variable cell type, tissue, and developmental contexts relevant to disease progression may yield critical information that would be lost by fine-tuning on each context separately. Furthermore, simultaneously learning about a broad range of diseases, cell types, tissues, and developmental stages may also yield a pre-fine-tuned model that could be directly used for in silico treatment analysis without the need to fine-tune to each setting separately. This would additionally allow one to test in silico perturbations for potential side effects in shifting towards a compendium of alternate diseases as opposed to the healthy state.

To address this, we developed a multi-task learning strategy to enable the model to learn context-specific representations of cell states from multiple cross-informative biologically-relevant tasks (Fig. 3a). We leveraged the CELLxGENE database^14^, which at the time of access contained ∼43 million singlecell transcriptomes with annotations across multiple biologically-informative features including, after further curation/consolidation, 71 cell types, 38 tissues, 68 diseases, and 3 developmental stages. Because the distribution across these classes was significantly imbalanced, we iteratively balanced each feature to generate a subsampled dataset that maintained the diversity of the original corpus (Extended Data Fig. 2a).

**Fig. 3.**
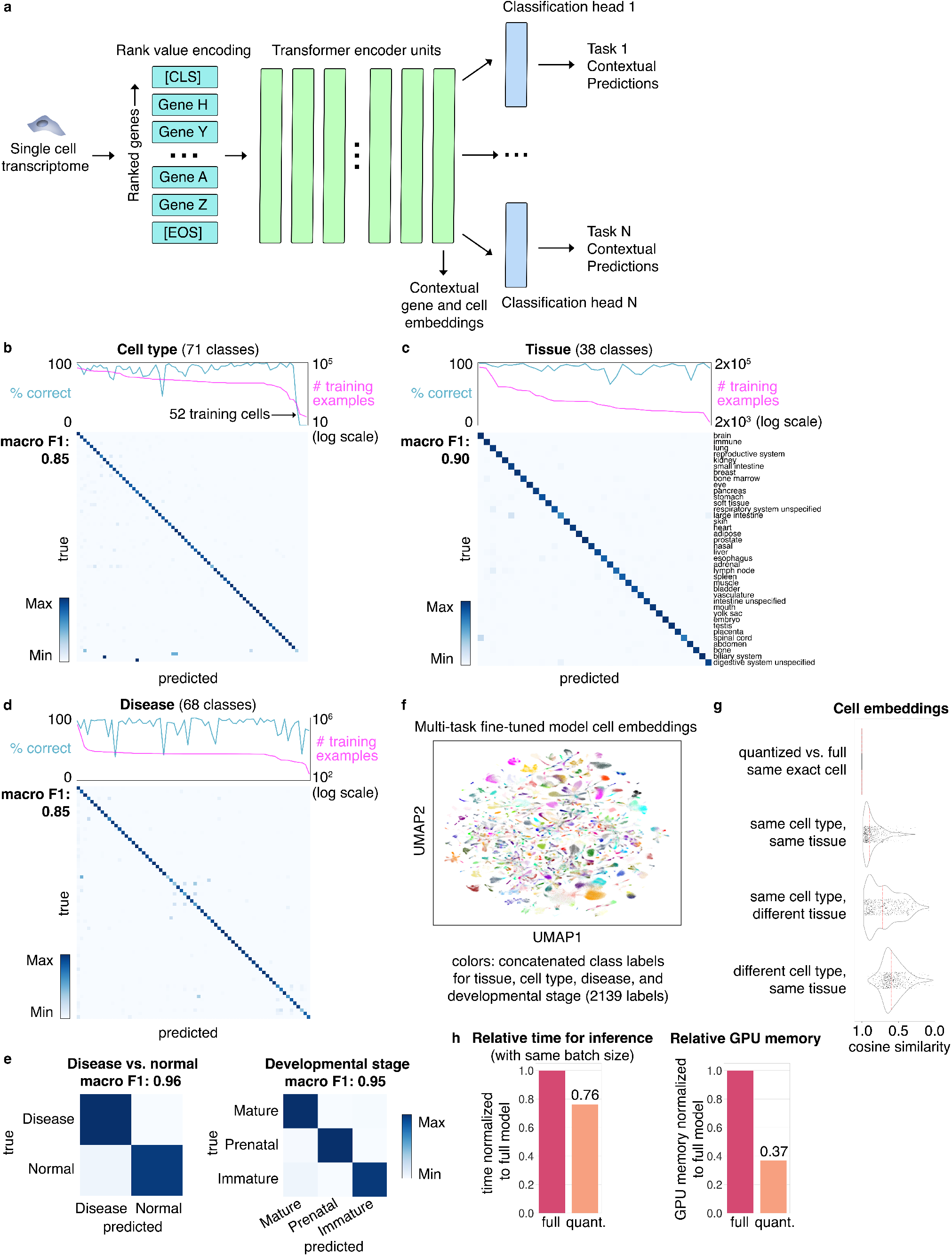
Multi-task learning yielded context-specific representations of cell states. **a**, Multi-task learning strategy starting from rank value encoding of each transcriptome with a CLS token for cell state classification followed by transformer encoder units with shared weights (12 layers in the case of GF-12L-95M-I4096) followed by a classification head for each task. The classification heads yield contextual predictions; contextual gene and cell embeddings can be extracted from each layer of the model. **b**, Cell type (71 classes), **c**, tissue (38 classes), **d**, disease (68 classes), **e**, disease vs. normal (2 classes), and developmental stage (3 classes) task confusion matrices and macro F1 scores for GF-12L-95M-I4096 jointly fine-tuned towards the aforementioned five tasks with the CELLxGENE corpus. Binary disease classification task was meant to instruct the model to understand that multiple diseases may exist for a given cell state that are all varying subtypes of dysfunction but altogether differ in a critical way from the normal state. Predictions (teal line) were robust to decreasing numbers of training examples (magenta line) down to an increasingly minimal number of examples (e.g. 52 training cells in the case of cell type). **f**, Second to last layer CLS cell embeddings from the multi-task fine-tuned GF-12L-95M-I4096 for a balanced subset of CELLxGENE with colors indicating concatenated class labels for the five tasks (total 2139 labels). **g**, Cosine similarity of CLS cell embeddings from the multi-task fine-tuned GF-12L-95M-I4096 of 3000 representative normal adult cells from a broad range of cell types and tissues from the CELLxGENE corpus. The top plot shows cosine similarity of embeddings generated from the 8-bit quantized vs. full model for the same exact cells. The below plots show cosine similarities of the full model’s embeddings for cells that are the same or different cell types and/or from the same or different tissues. **h**, Relative time for inference or relative GPU memory usage for the 8-bit quantized vs. full multi-task fine-tuned GF-12L-95M-I4096. Of note, time was quantified per the same batch size, but the actual time gains would be greater because the lower memory requirements of the model would allow larger batch sizes.

Jointly fine-tuning across these multiple biologically-informative tasks yielded a fine-tuned model with a validation macro F1 score of 0.85 for cell types, 0.90 for tissues, 0.85 for diseases, and 0.95 for developmental stages (Fig. 3b-e, Extended Data Fig. 2b, 3). The model was robust to decreasing amounts of input data per class, with performance dropping only with increasingly limited numbers of examples, such as cell type classes with 52 cells or less (Fig. 3b). Furthermore, many cases of label confusion were likely due to imprecise labeling, for example with the label of “digestive system unspecified” being classified by the model as “large intestine” or “small intestine” (Extended Data Fig. 2c). In the cases of the disease features, there were certainly some lowly represented diseases that were commonly misclassified by the model. However, in many cases of label confusion, the confused labels represented diseases with shared pathologies, such as frontotemporal dementia and amyotrophic lateral sclerosis, or Parkinson disease and Lewy body dementia (Extended Data Fig. 2d). The model was also able to apply CELLxGENE cell type labels to an external cross-tissue atlas^15^ (Extended Data Table 2, Extended Data Fig. 3).

The multi-task fine-tuning yielded context-specific representations of the cell states, defining within the embedding space a joint total of 2139 label combinations comprising cell type, tissue origin, developmental stage, and disease status of each individual cell (Fig. 3f). This defined embedding space may now be utilized as a reference for embedding new cells and to predict perturbations that shift between the represented states.

### Multi-task model quantization allowed resource-efficient in silico perturbation analysis

Because even inference with large models can be resource-intensive, we tested whether 8-bit quantization of the multi-task model could preserve the embedding space while increasing computational efficiency. Indeed, when testing normal adult cells from a broad range of tissues and cell types, the quantized embeddings mapped to nearly equivalent positions within the embedding space while reducing memory usage by 63% (Fig. 3g-h, Extended Data Fig. 4a). The time for inference was also 24% faster using the same batch size, though the maximum speed gains are much greater due to the ability to run larger amounts of data per batch given the significant reduction in memory usage.

Given that the quantized model embedding space preserved the context-specific biological representations of the cells, we tested whether the quantized model could be used for resource-efficient in silico perturbation analysis. As an example, we tested in silico deletion of genes from the transcriptional regulatory network database (TRRUST)^16^ in intestinal fibroblasts from patients with inflammatory bowel disease to determine the cosine shift towards the control intestinal fibroblast state and observed that the quantized model resulted in nearly equivalent shifts to the full model (Pearson correlation 0.9995) (Extended Data Fig. 4b). Thus, quantization of the multi-task model enabled resource-efficient embedding extraction and in silico perturbation analysis.

### Continual learning enabled domain-tuning for cancer states excluded from pretraining to boost predictions in colorectal cancer multi-task learning

The tumor microenvironment is an example of a disease setting affected by multicellular pathology where context-dependent gene network dysregulation in immune, stromal, and malignant cells influences tumor progression and the anti-tumor immune response^17^. Our multi-task learning strategy may be uniquely suited to modeling these highly context-specific states to determine candidate therapeutic targets in each cell type and tumor context that would promote anti-tumor immune responses. Yet, as discussed above, during pretraining we excluded malignant cells due to their propensity for gain of function mutations. These mutations would result in many genes with very different functions than what the model would observe in other contexts without accompanying genome sequencing to provide this information to the model. However, excluding cancer studies from the pretraining may result in the model having a lower baseline understanding of the gene network rewiring that occurs in malignancy.

Therefore, we performed domain-specific continual learning to tune Geneformer to the cancer domain by extending the pretraining with ∼14 million cells from cancer studies including matched healthy controls to provide this contrasting context to the model (Fig. 4a). We also included 1% of the non-cancer cells from Genecorpus103M to prevent catastrophic forgetting of the general knowledge of gene network dynamics learned by the model during the initial pretraining. We tested three different continual learning strategies and observed the lowest continual learning loss when the learning rate was rewarmed to the maximum learning rate used during the general pretraining and then decayed with a cosine schedule (Fig. 4b). When we used a learning rate that was 10% of the maximum used in the initial pretraining, there was minimal difference in the continual learning loss whether we left the learning rate constant or whether we rewarmed to that rate and decayed on a cosine schedule thereafter.

**Fig. 4.**
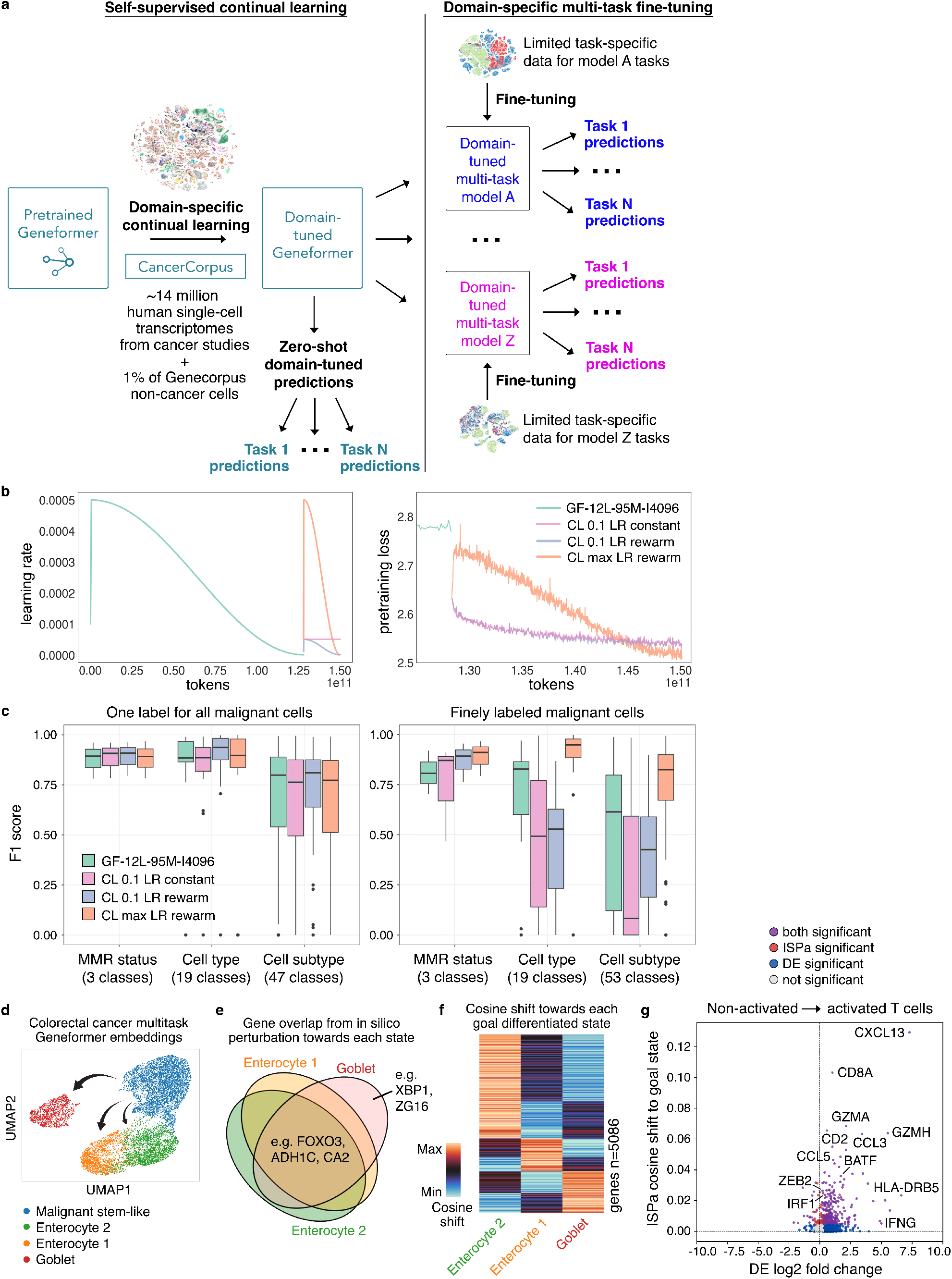
Domain-specific continual learning boosted predictions in domain-relevant multi-task learning. **a**, Domain-specific continual learning strategy starting from pretrained Geneformer and performing continual learning with ∼14 million human single-cell transcriptomes from cancer studies plus 1% of the Genecorpus non-cancer cells to prevent forgetting of baseline knowledge of network dynamics gained from the initial pretraining. The domain-tuned Geneformer can then perform zero-shot domain-tuned predictions or be transferred to downstream multi-task models for domain-specific fine-tuning. **b**, Learning rate and pretraining loss for each continual learning model. Teal indicates the initial pretraining of GF-12L-95M-I4096 prior to continual learning; the pretraining loss plot begins at the end of the initial pretraining to primarily display the continual learning phase. **c**, Multi-task fine-tuning results for each continual learning model compared to the pretrained GF-12L-95M-I4096 without continual learning. Models were fine-tuned using a colorectal cancer atlas17 to distinguish cell type, cell subtype, and MMR status of single-cell transcriptomes from MMRd or MMRp colorectal cancer specimens as well as within non-neoplastic colon epithelium from the same surgical resections, respectively. In models displayed in the left plot, all malignant cells were labeled as a single cell subtype; whereas in models displayed in the right plot, malignant cells were finely labeled as different malignant epithelial cell states such as stem-like or Goblet-like. **d**, UMAP of colorectal cancer multi-task Geneformer model embeddings for MMRd malignant transit-amplifying (TA)/stem-like epithelial cells and non-malignant differentiated epithelial cells of the Enterocyte 2, Enterocyte 1, and Goblet subtypes. Arrows indicate transitions modeled by the in silico perturbation analysis. **e**, Area-proportional Venn diagram of overlap of genes whose in silico activation significantly shifted the malignant TA/stem-like epithelial cells to each of the indicated non-malignant differentiated epithelial states (Wilcoxon with Benjamini-Hochberg (BH) correction, p<0.05 and greater shift than mean shift of all genes). **f**, Heatmap of cosine shifts towards each goal non-malignant differentiated epithelial state from the start state of malignant TA/stem-like epithelial cells in response to each gene’s in silico activation. Values are row-scaled. **g**, In silico treatment analysis to determine genes whose increased expression are predicted to induce non-activated T cells in MMRp tumors to shift towards the activated T cell state found in MMRd tumors with higher anti-tumor immune activation. ISPa=in silico perturbation, activation (Wilcoxon with BH correction, p<0.05 and greater shift than mean shift of all genes). DE=differential expression in goal end vs. start state (Wilcoxon with BH correction, p<0.05). ISPa positive hits significantly overlapped with differential expression positive hits (p<0.05, *X*^2^ test).

We then performed multi-task fine-tuning to learn context-specific representations of cell states within mismatch repair-deficient (MMRd) or -proficient (MMRp) colorectal cancer specimens as well as within nonneoplastic colon epithelium from the same surgical resections, respectively, using a colorectal cancer atlas^17^ (Fig. 4c, Extended Data Fig. 5a-c). Importantly, MMR status distinguishes patients with a favorable (MMRd) or poor (MMRp) rate of response to immunotherapy^18^. We fine-tuned the model to distinguish cells based on MMR status of the tumor, cell type classes, and finely labeled cell state classes and tested the ability of the model to predict these features in held-out patients.

When we binned all malignant cells into a single label of “malignant” in the cell subtype classes, the initial pretrained model was able to match the performance of the cancer-tuned models in distinguishing normal epithelial cells from the general category of malignant cells. However, the continually pretrained model that was rewarmed to the maximum learning rate used in the initial pretraining, and therefore most highly tuned to the cancer domain, outperformed the initial pretrained model when tasked to distinguish finely labeled malignant cells. Furthermore, the continual learning improved the ability of the model to distinguish the MMR status of each cancer, which is an important feature associated with differential immunotherapy response. Overall, the continual learning on cancer cells improved the ability of the model to distinguish the subtleties of malignant cells within the colorectal tumor microenvironment.

### In silico treatment analysis with multi-task colorectal cancer model predicted core effectors and regulators of malignant epithelial and activated T cells

We then leveraged the multi-task fine-tuned model for colorectal cancer to predict regulators that shift highly proliferative transit-amplifying/stem-like epithelial cancer cells in MMRd colorectal cancer to differentiated and thus likely less tumorigenic epithelial cell states found in normal epithelium, namely Enterocyte 1, Enterocyte 2, and Goblet cells (Fig. 4d, Extended Data Table 3). Genes whose in silico overexpression was predicted to shift away from the malignant stem-like state towards the differentiated states were significantly enriched for genes involved in key functions of intestinal epithelial cells, including nutrient transporters and microvillus structures, as well as genes related to cell polarity, which is known to get lost during cancer progression, promoting tumor growth^19^ (Fig. 4e-f, Extended Data Table 4). These genes included genes previously reported to inhibit tumor progression upon experimental overexpression such as CA2^20^ and ADH1C^21^. Furthermore, one of the top transcription factors whose in silico overexpression was predicted to shift towards the differentiated states was FOXO3, which was previously shown to suppress epithelial stem/progenitor cell states^22^. By contrast, one of the transcription factors that ranked specifically high in the shift to Goblet cells, but not Enterocytes, was XBP1, an endoplasmic reticulum stress transcription factor central to mucus-producing Goblet cells that was previously shown to be required to suppress intestinal tumorigenesis in a knockout mouse study^23^. The in silico perturbation analysis using the multi-task model therefore prioritized genes based on predicted shifts towards several distinct alternate end states.

Next, we leveraged the multi-task fine-tuned model for colorectal cancer to predict core effectors and central regulators of activated T cells found in the immunotherapy-responsive tumor microenvironment of MMRd tumors. Such genes could be interesting therapeutic targets to synthetically enhance the potency of adoptive cell therapies. We performed in silico treatment analysis to determine gene perturbations that shift the cell embeddings from an immunologically quiescent T cell state found in MMRp tumors to the activated T cell state specific to the immunotherapy-responsive MMRd colorectal cancer subtype (Fig. 4g, Extended Data Table 3). Overall, genes whose increased expression was predicted to shift T cells to the interferon gamma-producing, MHCII-positive T cell state selectively found in MMRd tumors^17^ were enriched for genes involved in type II interferon response, cytokine activity, inflammatory response, TNF-alpha signaling, and interleukin/STAT signaling (Extended Data Fig. 5d, Extended Data Table 5).

In silico perturbation analysis had higher concordance with a prior CRISPR activation screen^24^ in primary CD8+ T cells for interferon gamma-promoting perturbations compared to differential expression analysis (Extended Data Fig. 5e, Extended Data Table 3). Among the top 25 significant genes were CXCL13 (top hit), which is a central gene in tumor-reactive T cells across multiple tumor types^25^, granzymes and granulysin that are important for T cell cytotoxicity, and components of the T cell receptor. The top 3 significant transcription factors^26^ were BATF, which counters T cell exhaustion and promotes effector function^27^, ZEB2, which promotes terminal differentiation of CD8+ effector and memory T cells^28^, and IRF1, a core driver of interferon stimulated genes and Th1 differentiation programs in CD4+ T cells^29^. Thus, the multi-task learning approach enabled predictions of core genes of activated T cells in immunotherapy-responsive tumor microenvironments, including known targets previously shown to improve T cell function.

## Discussion

In sum, we developed a quantized multi-task learning strategy to yield context-specific representations of gene network dynamics that can be leveraged to make contextual predictions of key network regulators and candidate therapeutic targets for human disease. Expanding the pretraining corpus from Geneformer’s initial ∼30 million^6^ to now ∼95 million cells as well as increasing the model parameters and input context size to 4096 improved zero-shot performance on diverse downstream tasks relevant to gene network biology, chromatin dynamics, and disease modeling. As the quantity and diversity of available single-cell transcriptomic data continues to grow, future larger models pretrained on even larger-scale corpuses may open opportunities to achieve meaningful predictions in even more elusive tasks as zero or few-shot learners.

For domains underrepresented in the pretraining corpus, such as the case of the cancer domain presented here, continual learning may serve as a viable method to tune the model for predictions in these specific domains. Furthermore, multi-task fine-tuning allowed the model to jointly learn diverse biological dimensions critical to defining a cell’s state, which may be particularly important in modeling developmental dynamics and diseases with multicellular pathology such as the tumor-immune microenvironment.

Finally, we demonstrate that model quantization with QLoRA is an effective method for resource-efficient finetuning, embedding extraction, and in silico perturbation for biological applications. As model size and data grow while available GPU resources remain a limitation, approaches for efficient fine-tuning and inference will be critical to ensure widespread access to models for biological discoveries that have the potential to impact human health.

Overall, quantized multi-task learning enables resource-efficient context-specific modeling in gene network biology to yield contextual predictions of key network regulators and candidate therapeutic targets for human disease.

## Methods

Complete methods available in Extended Data.

## Data Availability

Genecorpus-103M will be available on Hugging Face Dataset Hub.

## Code Availability

The pretrained Geneformer-95M models, cancer-tuned model, and multi-task fine-tuned model and related code are available on Hugging Face Model Hub at https://huggingface.co/ctheodoris/Geneformer.

## Acknowledgements

We thank Jack Rae and the Theodoris Lab members for helpful scientific discussions and Argonne National Laboratory for providing GPU resources for experimentation. CVT was supported by grants from the National Institutes of Health (DP5OD036170), Burroughs Wellcome Fund Career Award for Medical Scientists (1022136.01), Biswas Foundation, and Milken Institute. KP was supported by NIH/NCI grant R00CA259511, the NIH/NCI Cancer Cell Map Initiative, the Parker Institute for Cancer Immunotherapy, funds from the CRISPR Cure for Cancer Initiative, the UCSF Program for Breakthrough Biomedical Research, Biswas Foundation, and Milken Institute. HC was supported by the National Science Foundation Graduate Research Fellowship (2034836). TNN and RKM were supported by the United States Department of Energy Office of Science-Advanced Scientific Computing Research Program (Contract No. DEAC02-06CH11357).

## Author Contributions

HC and MSV developed the models and designed/performed computational analyses. HC assembled the pretraining corpuses and developed the continual learning method. MSV developed the multitask learning method and quantization strategy. JGO contributed to model pretraining and corpus assembly. SVM contributed to benchmarking. TNN and RKM advised on distributed training and contributed to the continual learning method. KP and CVT designed analyses and supervised the work. HC, MSV, KP, and CVT wrote the manuscript. All authors edited the manuscript.

## Extended Data

### Extended Methods

#### Assembly and rank value encoding of transcriptomes in Genecorpus-103M

##### Assembly and uniform processing of single-cell transcriptomes

We assembled a large-scale pretraining corpus, Genecorpus-103M, comprising ∼103 million human single-cell transcriptomes (post-filtering as described below) from a broad range of tissues from 4,610 publicly available datasets (Fig. 1b, Extended Data Table 1). Importantly, DOIs were cross-referenced between all studies to ensure datasets were unique to avoid inclusion of duplicated cells within the corpus. Of note, there are significant duplications of datasets across public databases so the total number of unique cells would be highly overestimated if this procedure were not performed (Extended Data Fig. 1a).

Publicly available datasets containing raw counts were collected from National Center for Biotechnology Information (NCBI) Gene Expression Omnibus (GEO), NCBI Sequence Read Archive (SRA), CELLxGENE, Human Cell Atlas, European Molecular Biology Laboratory-European Bioinformatics Institute (EMBL-EBI) Single Cell Expression Atlas, Broad Institute Single Cell Portal, Brotman Baty Institute (BBI)-Allen Single Cell Atlases, Tumor Immune Single-cell Hub (TISCH) (excluding malignant cells), Panglao Database, 10x Genomics, University of California, Santa Cruz Cell Browser, European Genome-phenome Archive, Synapse, Riken, Zenodo, National Institutes of Health (NIH) Figshare Archive, NCBI dbGap, Refine.bio, China National GeneBank Sequence Archive, Mendeley Data, and individual communication with authors of the original studies (Extended Data Table 1). Additional resources for collecting information about suitable studies included Entrez Direct tools and the dataset summary from Svensson et al., Database 2020^30^. Tools utilized in conversion of data to uniform files included loompy, scanpy, anndata, scipy, numpy, pandas, Cellranger, and LoomExperiment. Gene annotation data was retrieved from Ensembl, NCBI, and HGNC (2023-11-01) databases and additionally queried through MyGene^31^. Raw and unfiltered data files were processed to remove empty droplets and debris using STAR version 2.7.8a^32^ with the CellRanger2.2 (run mode –soloCellFiltered). Datasets were additionally filtered to retain cells that contained a minimum of seven detected Ensembl-annotated protein coding genes given that the 15% masking used for the pretraining learning objective would not reliably mask a gene in cells with fewer detected genes. Studies were annotated as one or more of the 52 consolidated organs as listed in Fig. 1b.

##### Rank value encoding of single-cell transcriptomes

Each transcriptome was presented to the model as a rank value encoding as previously described^6^. The rank value encodings are a nonparametric representation of the transcriptome that takes advantage of the many observations of the gene’s expression across the entire Genecorpus to prioritize genes that distinguish cell state. Specifically, this method will deprioritize ubiquitously highly-expressed housekeeping genes by normalizing them to a lower rank. Conversely, genes such as transcription factors that may be lowly expressed when they are expressed but highly distinguish cell state will move to a higher rank within the encoding. Furthermore, this rank-based approach may be more robust against technical artifacts that may systematically bias the absolute transcript counts value while the overall relative ranking of genes within each cell remains more stable.

The rank value encodings were constructed as previously described^6^. The scaling factor for each gene was derived from the non-zero median value of expression of each detected gene across all cells in the pretraining corpus passing quality filtering that were sequenced on droplet-based platforms, excluding cells with high mutational burdens such as malignant cells and immortalized cell lines. After scaling the expression of each gene, the genes were ordered by the rank of their scaled expression in that specific cell. The rank value encoding for each single-cell transcriptome was then tokenized on the basis of a vocabulary of 20,271 protein coding genes detected within the pretraining corpus. The vocabulary also included four special tokens: a padding, masking, CLS (classification), and EOS (end of state) token, for a total vocabulary size of 20,275. A CLS and EOS token were added to the beginning and end of each rank value encoding, respectively. The tokenized dataset was stored within the Hugging Face Datasets structure, which is based on the Apache Arrow format that allows processing of large datasets with zero-copy reads without memory constraints.

Of note, this strategy is also space-efficient as the genes are stored as ranked tokens as opposed to the exact transcript values, and we only store genes detected within each cell rather than the full sparse dataset that includes all of the undetected genes. This also prevents wasting computation on zeros, as the model learns from the absence of genes from a rank value encoding without having to explicitly instruct the model that they have zero expression. This is analogous, for example, to how natural language models learn that a statement may have “positive” meaning based on the absence of “negative” words, without needing to present the remainder of the absent words from the natural language dictionary at the end of every sentence to explicitly instruct the model they are not present.

#### Geneformer architecture and pretraining

##### Geneformer architecture

In the initial pretraining phase, Geneformer was pretrained with ∼95 million (94,222,639) single-cell transcriptomes from Genecorpus-103M excluding cells with high mutational burdens such as malignant cells and immortalized cell lines. Geneformer-95M is composed of 12 or 20 transformer encoder units, each composed of a self-attention layer and feed forward neural network with the following parameters, respectively: input size of 4096 (fully represents 93% of Genecorpus-103M), 512 or 896 embedding dimensions, 8 or 14 attention heads per layer, and a feed forward size of 1024 or 1792. Further parameters are as follows: nonlinear activation function: rectified linear unit (ReLU); dropout probability for for all fully connected layers: 0.02; dropout ratio for attention probabilities: 0.02; standard deviation of the initializer for weight matrices: 0.02; epsilon for layer normalization layers: 1e-12. We additionally pretrained a Geneformer-95M version with input size of 2048 that otherwise had the same parameters as the 12 layer model above. We compared these Geneformer-95M models to the previous Geneformer models pretrained on ∼30 million transcriptomes that were either 6 layers as previously described^6^ or 12 layers with parameters as the 12 layer model above except with input size of 2048. Modeling was implemented in pytorch and using the Hugging Face Transformers library for model configuration, data loading, and training.

##### Geneformer pretraining and performance optimization

Pretraining was accomplished using a masked learning objective, which has been shown in other informational fields^3-4^ to improve generalizability of the foundational knowledge learned during pretraining for a wide range of downstream fine-tuning objectives. During pretraining, 15% of the genes within each transcriptome were masked, and the model was trained to predict which gene should be within each masked position in that specific cell state using the context of the remaining unmasked genes. A major strength of this approach is that it is entirely self-supervised and can be accomplished on completely unlabeled data, which allows the inclusion of large amounts of training data without being restricted to samples with accompanying labels. Pretraining hyperparameters were optimized to the following final values: max learning rate: 20 layer (L)-Input size (I) 4096: 2.5e-4, 12L-I4096: 5e-4, 12L-I2048: 1e-4; learning scheduler: cosine with warmup; optimizer: Adam with weight decay fix; warmup steps: 920 for 20 layer model and 5,000 for 12 layer models; weight decay: 0.001; batch size: 1 with gradient accumulation 12 for 20 layer model and 4 for 12 layer models. Tensorboard was used for experimentation tracking, and each model was pretrained for 3 epochs.

As the input size of 4096 is considerably large for a full dense self-attention model (for example, BERT^3-4^ input size of 512) and transformers have a quadratic memory and time complexity *O*(*L*^2^) with respect to input size, we implemented measures to optimize efficiency of large-scale pretraining. The trainer from the Hugging Face Transformers library was used for pretraining with the substitution of a custom tokenizer to implement dynamic, length-grouped padding, which minimized computation on padding and achieved a 29.4x speedup in pretraining. This process takes a randomly sampled megabatch and then orders minibatches by their length in descending order (to ensure that any memory constraints are encountered earlier). Minibatches are then dynamically padded, minimizing the computation wasted on padding due to being length-grouped. We also implemented distributed GPU training algorithms^33-34^ to allow efficient pretraining on the large-scale dataset using Deepspeed, which partitions parameters, gradients, and optimizer states across available GPUs, offloads processing/memory as possible to CPU to allow more to fit on GPU, and reduces memory fragmentation by ensuring long and short term memory allocations do not mix. Overall, pretraining for the 20L-I4096, 12L-I4096, and 12L-I2048 Geneformer-95M models was achieved in approximately 134, 44, or 29 hours, respectively, each distributed across 8 Nvidia H100 80GB GPUs.

#### Zero-shot learning evaluation for the pretrained models

The five pretrained models (GF-6L-30M-I2048, GF-12L-30M-I2048, GF-12L-95M-I2048, GF-12L-95M-I4096, GF-20L-95M-I4096) (GF=Geneformer, L=Layers, M=Million cells, I=Input size) were evaluated on gene classification tasks including disease genes (dosage sensitive vs. insensitive transcription factors), chromatin dynamics (bivalent vs. lys4-only methylated genes from 56 highly conserved loci), network dynamics (central vs. peripheral genes within the defined NOTCH1-dependent gene network), and gene regulation (long range vs. short range transcription factors) as previously described^6^. The example cells were as previously described^6^, except that in the chromatin dynamics and network dynamics tasks the number of example cells was reduced to increase the difficulty of the task. The example cells, as previously described^6^ for each task were: disease genes: 10,000 random cells from the respective pretraining corpus of each model; chromatin dynamics: 10,000 embryonic stem cells^35^; network dynamics: 2,000 normal endothelial cells^36^; and gene regulation: ∼34,000 cells from an iPSC to cardiomyocyte differentiation^37^. Training was performed on 80% of the genes in each task by freezing all model layers and training a classification head on the zero-shot gene embeddings for a single epoch. 25 hyperparameter trials for the classification on the zero-shot embeddings were employed to ensure equitable comparison between the models with search ranges of maximum learning rate 1e-6 to 1e-3, weight decay 0.0 to 0.3, and seed 0 to 100 (to preclude the chance that a randomly selected seed would be advantageous for classification of zero-shot embeddings of one model over another). Performance was evaluated on classification of the zero-shot embeddings of the held-out genes.

Zero-shot cell embeddings were extracted from the second to last layer CLS token from GF-12L-95M-I4096 for 779,905 representative cells from CELLxGENE^14^ balanced across cell types, tissues, diseases, and developmental stages as discussed in the Multi-task learning section below and colored by consolidated labels for those cell attributes as discussed below.

#### Model quantization

Quantization of the GF-20L-95M-I4096 model was performed using 4-bit QLoRA^13^. For Fig. 2a, the four tested task types were as described in the *Zero-shot learning evaluations* section above, with 80% of the genes being used for training. Performance was evaluated on classification of the held-out genes. QLoRA models were trained with rank of 16 and alpha of 32. Three maximum learning rates were tested (0.0005, 0.0001, and 0.00005), all with a warmup ratio of 1% and learning rate decay on a cosine schedule, with either 0, 7, or 14 layers being frozen from fine-tuning. Full fine-tuning was performed for the GF-20L-95M-I4096 model with 25 trials as described in the *Zero-shot learning evaluations* section above but instead of freezing all layers, only 0, 7, or 14 layers were frozen, allowing the remainder to be fine-tuned. Zero-shot learning with the GF-20L-95M-I4096 model was performed as described in the *Zero-shot learning evaluations* section above. All models were trained for a single epoch with batch size of 1 and gradient accumulation of 12.

For the few-shot learning application in Fig. 2c, the disease gene task was performed as described above but with only 100 randomly subsampled example cells. All layers were allowed to be fine-tuned for both QLoRA and full fine-tuning models (0 layers were frozen). QLoRA rank was either 16, 32, 64, or 128, and alpha was twice the rank. For both QLoRA and full fine-tuning models, the maximum learning rate was 0.0001, decayed on a cosine schedule with warmup ratio of 1%. All models were trained for a single epoch with batch size of 1. For each model, the training was repeated with three different seeds. 80% of the genes were used for training and performance was evaluated on classification of the held-out genes.

The multi-task model fine-tuned on cells from the CELLxGENE corpus was quantized from full precision (FP32) to 8-bit precision for inference. Specifically, the quantization configuration loaded the model in 8-bit precision without employing low-rank matrices A and B for fine-tuning. The full precision weights were directly converted to a lower precision without additional low-rank factorization, such that the original architecture and weights remained intact during the quantization process.

#### Domain-specific continual learning

During the initial pretraining phase, cancer studies were excluded due to the propensity for malignant cells to harbor gain of function mutations that would result in many genes with very different functions than what the model would observe in other contexts without accompanying genome sequencing to provide this information to the model. However, because this may result in the model having a lower baseline understanding of the gene network rewiring that occurs in malignancy, we performed domain-specific continual learning to tune Geneformer to the cancer domain. We extended the pretraining of the 12L-I4096 Geneformer model with ∼14 million (13,628,457) cells from cancer studies including both malignant cells and non-tumor cells in the tumor microenvironment and matched healthy controls to provide this contrasting context to the model. We also included 1% of the non-cancer cells from Genecorpus-103M to prevent catastrophic forgetting of the general knowledge of gene network dynamics learned by the model during the initial pretraining. We tested three different continual learning strategies: one where we rewarmed (warmup ratio 1%) to the maximum of the learning rate from the initial pretraining and decayed the learning rate on a cosine schedule (“CL max LR rewarm”), one where we rewarmed (warmup ratio 1%) to 10% of the maximum of the learning rate from the initial pretraining and decayed the learning rate on a cosine schedule (“CL 0.1 LR rewarm”), and one where we continued pretraining at a constant learning rate that was 10% of the maximum used for the initial pretraining (“CL 0.1 LR constant”). The other hyperparameters were the same as the initial pretraining for the 12L-I4096 model except that the models were continually trained for only 1 epoch.

#### Multi-task learning

##### Multi-task learning on CELLxGENE corpus

The multi-task learning architecture was composed of shared weights for all tasks within the trunk of the model initialized from the pretrained GF-12L-95M-I4096 model with a classification head on top for each task. Weights were updated during multi-task fine-tuning based on weighted task-specific losses, where each task’s cross-entropy loss contribution was individually scaled depending on its importance or complexity. The five tasks represented biologically meaningful attributes of cells annotated within the CELLxGENE^14^ corpus (accessed 4/12/2024) including cell type (consolidated to 71 classes), tissues (consolidated to 38 classes), diseases (consolidated to 68 classes), developmental stages (consolidated to 3 classes), and a binary indicator of whether the cell was annotated to be sampled from a healthy patient or one with one of the annotated diseases. The cell types represented the “cell subclass” attribute annotated by CELLxGENE. The labels for the cell types, tissues, and diseases were consolidated to reduce the occurrence of duplicate or imprecise labels or labels that occurred at varying levels of hierarchy within the relevant ontology. The developmental stages were consolidated to the CELLxGENE annotated labels of prenatal, immature (0-12 years of age), and mature (13+ years of age) as there was a substantial percentage of cells that were only labeled at this level of hierarchy (e.g. “mature”) so this consolidation was required to bring all labels to the same level of hierarchy. Cells without labels in all of the five tasks were excluded.

Of the ∼43 million cells in CELLxGENE at the time of access, we iteratively downsampled the dataset to better balance the classes in each of the above tasks. First, up to 5000 cells from each cell type were randomly selected. Then, cells were added to increase the representation of each of the diseases up to 5000 cells by randomly sampling from the underrepresented classes. Then, cells were added to increase the representation of each of the tissues up to 5000 cells by randomly sampling from the underrepresented classes. Then, cells were added to increase the representation of each of the developmental stages up to 100,000 cells by randomly sampling from the underrepresented classes. (This subsampled dataset comprised the 779,905 cells that were utilized for extracting zero-shot embeddings for GF-12L-95M-I4096 as discussed in the *Zero-shot learning evaluations* section above). For the held-out validation set, the same procedure was followed except that up to 1000 cells were randomly selected for each task class. For classes with less than 100 cells remaining in the CELLxGENE corpus after preparing the training dataset, we re-assigned 5% of the data for that class to use for validation, with only the remaining 95% being used for training. Ultimately, the training set comprised 776,709 cells and the validation set comprised 131,604 with no overlap between the dataset splits.

The multi-task model was then trained with the following optimal hyperparameters selected from 35 trials: max learning rate: 0.000355; learning scheduler: cosine with warmup; optimizer: Adam with weight decay fix; warmup steps: 600; weight decay: 0.05; batch size: 4; task weights: consolidated_cell_subclass: 0.3, consolidated_tissue: 0.2, disease_binary: 0.1, developmental_stage: 0.1, disease: 0.3; frozen layers: 10; epoch: 1. Out of sample performance was tested on the validation dataset described above. Cell embeddings were extracted from the second to last layer CLS token for the 779,905 cells in the subsampled dataset described above and colored by concatenated labels for each task for a total of 2139 labels.

Cell type label transfer was tested using an external cross-tissue atlas^15^ (not in CELLxGENE). First, a cell type label translation dictionary was manually curated to translate between the cell type labels annotated by the authors of the cross-tissue atlas and the cell type labels used by CELLxGENE. Of note, in many cases, there was not a direct corollary to the cross-tissue atlas labels within the CELLxGENE labels, the labels occurred at varying levels of hierarchy, and there was not a 1:1 mapping between label sets. The translation dictionary is provided in Extended Data Table 2. The multi-task model fine-tuned on the CELLxGENE data as described above was then used for inference, without further training, to predict CELLxGENE cell type labels for the cells within the external cross-tissue atlas.

##### Domain-tuned multi-task learning on a colorectal cancer atlas

Multi-task learning was performed leveraging a colorectal cancer atlas^17^ (GSE178341) using the pretrained GF-12L-95M-I4096 model (with or without cancer domain-tuning via the three methods described above in the Domain-specific continual learning section) with a classification head on top for each of the three tasks as follows: MMR status (3 classes: MMRd, MMRp, non-malignant), cell type (19 classes, consistent with “ClusterMidway” annotation from GSE178341), and cell subtype. The cell subtype consisted of 47 or 53 classes, depending on whether all malignant cells were labeled as one class or as finer subclasses, respectively. Patients from the colorectal cancer dataset^17^ were split into training (80%, n=50), validation (10%, n=6), and test (10%, n=6) sets by individual patients, balancing attributes including sex, MMR status, histological grade, and lymph node status. Hyperparameters were optimized over 25 trials and the model with the best performance on the validation set was then tested for generalizability on the held-out patients in the test set. The reported results are the performance on the held-out patients in the test set. The final best hyperparameters were: max learning rate: 0.000122859; learning scheduler: cosine with warmup; optimizer: Adam with weight decay fix; warmup steps: 680; weight decay: 0.036751018; dropout rate: 0.341148666; batch size: 3; task weights: 0.234978 (MMR status), 0.392293 (cell type), 0.370729 (cell subtype); frozen layers: 2; epoch: 1.

#### In silico treatment analysis

##### Inflammatory bowel disease in silico treatment analysis

In silico treatment analysis was performed as previously described^6^ with the multi-task model fine-tuned on the CELLxGENE corpus, with or without quantization (as described above in Model quantization). We tested in silico deletion of genes within the transcriptional regulatory network database (TRRUST)^16^ expressed in at least 20% of the start cells or goal end cells and measured the cosine shifts from the start population of fibroblasts from pediatric patients with Crohn disease to the goal end population of fibroblasts from age-matched control patients^38^.

##### Colorectal cancer in silico treatment analysis: T cells

In silico treatment analysis was performed with the multi-task fine-tuned model for colorectal cancer as previously described^6^ to determine genes whose increased expression would shift target cells within the tumor microenvironment to immune activating states. We tested in silico perturbations to shift CD8 T cells from an interferon gamma (IFNg) negative, non-activated state found in MMRp samples towards an IFNg positive, activated state observed in MMRd samples that are more immune-active. Cell states were as defined by the original authors, namely pTNI06 high versus low for activated versus non-activated T cells^17^.

We tested in silico activation of each gene that was detected in at least 20% of the start cells or goal end cells to determine candidates that were expressed in T cells that may be more viable to activate therapeutically. The goal end state was defined by the mean last layer CLS embedding position of a random subsample of 5000 cells from the goal end cell state. We tested in silico activation of each gene in a randomly subsampled subset of 5000 start state cells, where activation was modeled by moving the overexpressed genes to the first position in the rank value encoding after the CLS token.

We determined the genes whose activation in the start state cells significantly shifted the embeddings towards the goal end state embedding position within the 512-dimensional embedding space. Cosine similarities were quantified using the last layer CLS token embeddings for the original and in silico perturbed cells (last layer prior to the classification heads). Hits were defined as the genes whose in silico activation significantly shifted the cells from their original embedding position to the goal embedding position by cosine similarity as compared to the random distribution of random gene activation (p<0.05, Wilcoxon rank sum test, BH-corrected).

Gene set enrichment analysis was performed using the GSEApy^39^ implementation of Enrichr^40^ with the enrichment set being the genes whose in silico activation statistically significantly shifted the cells towards the goal state more than the average shift and the background set being all genes detected in at least 20% of cells. Enrichment sets were compared to the MSigDB Hallmark 2020, Kegg Human 2021, and Gene Ontology Biological Processes gene sets. For differential expression, raw counts were first preprocessed by count depth scaling with normalization of total counts to 10,000 followed by log1p transformation and scaling to a max value of 10 using Scanpy^41^. Differentially expressed genes were identified by comparing cells in the goal end state to the start cell state with Scanpy’s rank_genes_groups using Wilcoxon with BH multiple hypothesis correction.

To compare results with a prior CRISPR activation (CRISPRa) screen^24^ for IFNg-promoting perturbations in CD8+ T cells, CRISPRa hits annotated by the original authors were extracted from the referenced publication’s supplementary Table 2 with the following filters: “Screen_Version”=“Primary”, “CRISPRa_or_i”=“CRISPRa”, “CD4_or_CD8”=“CD8”, and “Cytokine”=“IFNg”. Positive hits were defined as those with the label “Positive Hit” in the column “Hit_Type”, while all other genes were considered not positive hits. These CRISPRa hits vs. not hits were compared with in silico perturbation-activation (ISPa) hits vs. not hits and differential expression hits vs. not hits. ISPa and differential expression analysis are described above. The ISPa start state was IFNg negative, non-activated state observed in the MMRp tumors and the goal end state was IFNg positive, activated T cells observed in the MMRd tumors. ISPa was performed with genes detected in at least 20% of the start cells or goal end cells to determine candidates that were expressed in T cells that may be more viable to activate therapeutically. For comparison with the CRISPRa results, ISPa positive hits were genes whose in silico activation in the non-activated T cell start state caused a statistically significant greater shift towards the goal end activated T cell state compared to the random distribution, whereas ISPa not positive hits were all other ISPa tested genes. Differential expression positive hits were those ISPa tested genes that were statistically significantly increased in expression in the goal end activated T cell state compared to non-activated T cell start state, whereas differential expression not positive hits were all other ISPa tested genes. Accuracy of concordance was calculated as (TP + TN) / (TP + TN + FP + FN) where T=true, F=false, P=positive, and N=negative for the CRISPRa positive hits vs. other genes compared to either the ISPa positive hits vs. other genes or the differential expression positive hits vs. other genes.

##### Colorectal cancer in silico treatment analysis: malignant epithelial cells

In silico treatment analysis was performed with the multi-task fine-tuned model for colorectal cancer as previously described^6^ and as detailed above, in this case to determine genes whose increased expression would shift stem-like malignant epithelial cells to one of three differentiated non-malignant epithelial cell states within the colon. Cell states were as defined by the original authors for transit-amplifying/stem-like malignant epithelial cells (cE01) and the three differentiated non-malignant epithelial cell states: enterocyte 1 (cE04), enterocyte 2 (cE05), and goblet (cE08)^17^. As discussed above, we tested in silico activation of each gene that was detected in at least 20% of the start cells or goal end cells and defined the goal end states by the mean last layer CLS embedding position of a random subsample of 5000 cells from each goal end state. In silico activation of each gene was tested in a randomly subsampled subset of 5000 start state cells. The cosine shift for in silico activation of each gene was compared to the random distribution of random gene activation (p<0.05, Wilcoxon rank sum test, BH-corrected). Gene set enrichment analysis was performed with ToppGene. The top 100 genes shifting from stem-like malignant to non-malignant epithelial cell states were defined as genes whose in silico activation led to a statistically significant shift towards all of the three differentiated states, ordered by the average shift to the three states. The background gene set was all genes tested by in silico activation.

**Extended Data Table 1** Dataset composition of Genecorpus-103M.

**Extended Data Table 2** Cell type label translation dictionary used to score correct vs. incorrect label transfer between the external cross-tissue atlas author labels and the closest analogous CELLxGENE labels.

**Extended Data Table 3** *Sheet 1*: Description of other sheets. *Sheet 2-4*: Genes whose in silico activation in malignant stem-like epithelial cells in MMRd tumors were predicted to significantly shift the cell state towards one of three non-malignant differentiated epithelial cells (Enterocyte 2, Enterocyte 1, or Goblet cells). *Sheet 5*: Genes whose in silico activation in non-activated T cells in MMRp tumors were predicted to significantly shift the cell state towards the activated T cell state found in MMRd tumors with higher anti-tumor immune activation. *Sheet 6*: Accuracy of concordance between T cell in silico activation and CRISPR activation screen in primary CD8+ T cells for interferon gamma production.

**Extended Data Table 4** Gene set enrichments of the top 100 genes whose in silico activation in malignant stem-like epithelial cells in MMRd tumors were predicted to significantly shift the cell state towards the three subtypes of non-malignant differentiated epithelial cells (Enterocyte 2, Enterocyte 1, or Goblet cells). The top 100 genes were ranked based on the largest average shift across all three non-malignant differentiated epithelial states.

**Extended Data Table 5** Gene set enrichments of genes whose in silico activation in non-activated T cells in MMRp tumors were predicted to significantly shift the cell state towards the activated T cell state found in MMRd tumors with higher anti-tumor immune activation.

**Extended Data Fig. 1.**
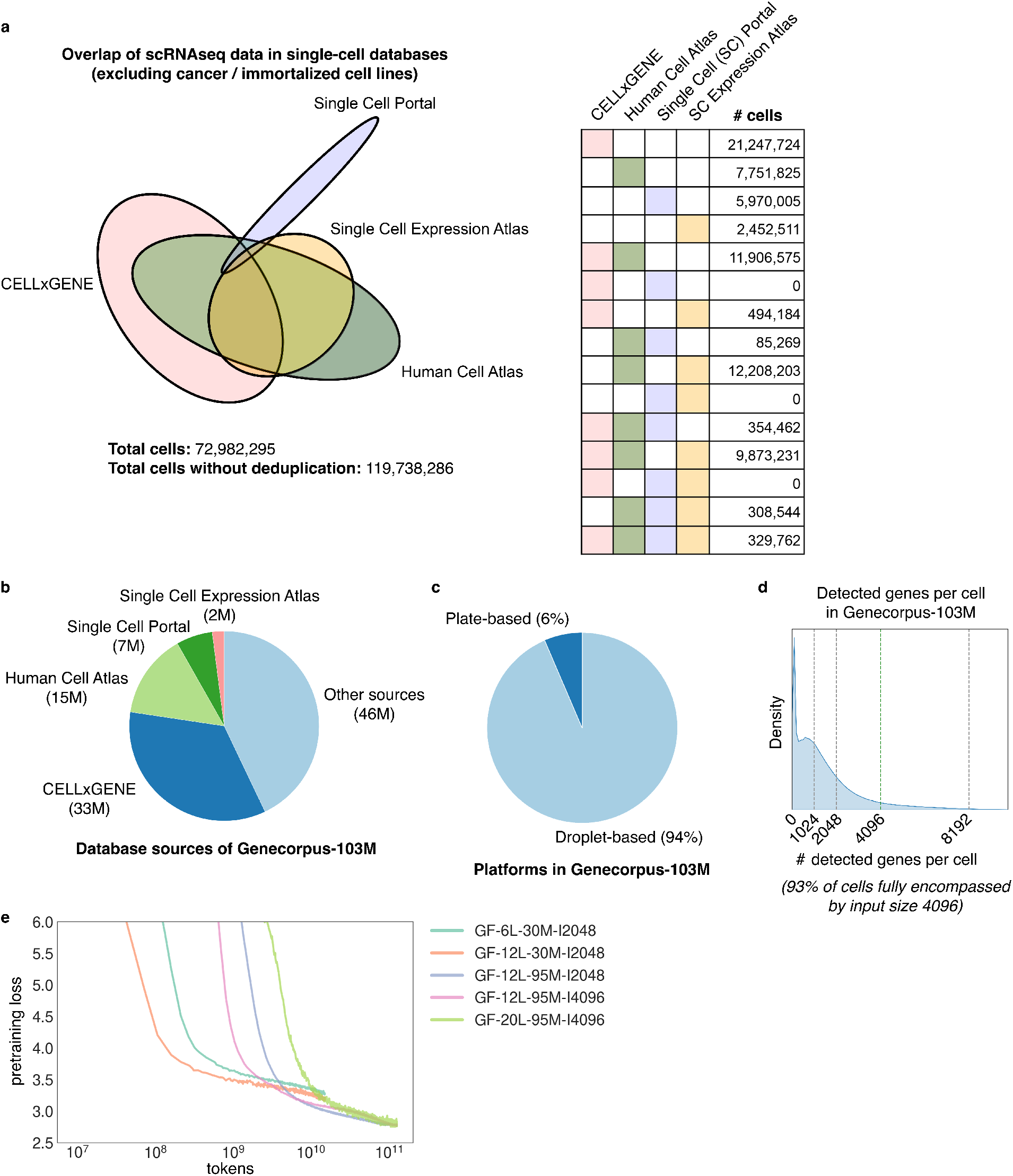
Pretraining corpus attributes. **a**, Overlap of single-cell transcriptome (scRNAseq) data from single-cell databases (excluding cancer / immortalized cell lines). Without deduplication by DOI, total cells are significantly overestimated. **b**, Database sources of Genecorpus-103M. **c**, Droplet-based vs. plate-based platform composition of Genecorpus-103M. **d**, Number of detected genes per cell in Genecorpus-103M. Input size of 4096 fully encompasses 93% of cells. **e**, Pretraining loss for each pretrained model (GF=Geneformer, L=Layers, M=Million cells, I=Input size) per number of tokens trained. Plot represents 3 epochs.

**Extended Data Fig. 2.**
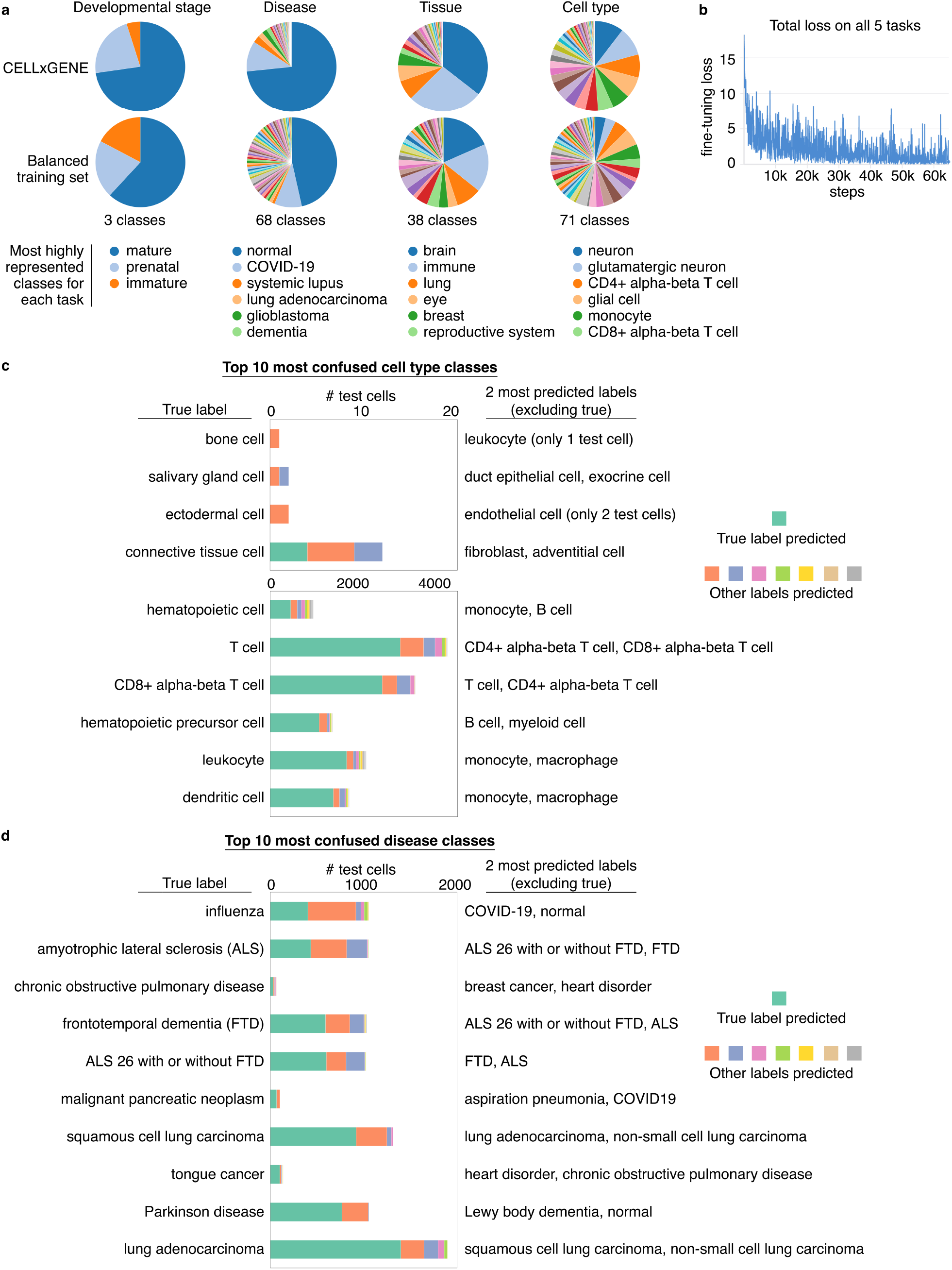
Multi-task learning dataset composition and predictions. **a**, Composition of original CELLxGENE corpus compared to the balanced training set used for multi-task learning for the four tasks of developmental stage, disease, tissue, and cell type. **b**, Total fine-tuning loss on all five tasks of cell type (71 classes), tissue (38 classes), disease (68 classes), disease vs. normal (2 classes), and developmental stage (3 classes). Plot represents 1 epoch. **c**, Top 10 most confused cell type classes and the top two most predicted labels excluding true (true was generally the top predicted label even in these top confused classes, as indicated by teal bar stacks). Alternate predicted labels were often cell types within the same overarching category (e.g. connective tissue cell and fibroblast, T cell and CD4+ alpha-beta T cell, dendritic cell and monocyte). **d**, Top 10 most confused disease classes and the top two most predicted labels excluding true (true was generally the top predicted label even in these top confused classes, as indicated by teal bar stacks). Alternate predicted labels were often diseases with shared pathologies (e.g. ALS and FTD; Parkinson and Lewy body dementia; influenza and COVID-19).

**Extended Data Fig. 3.**
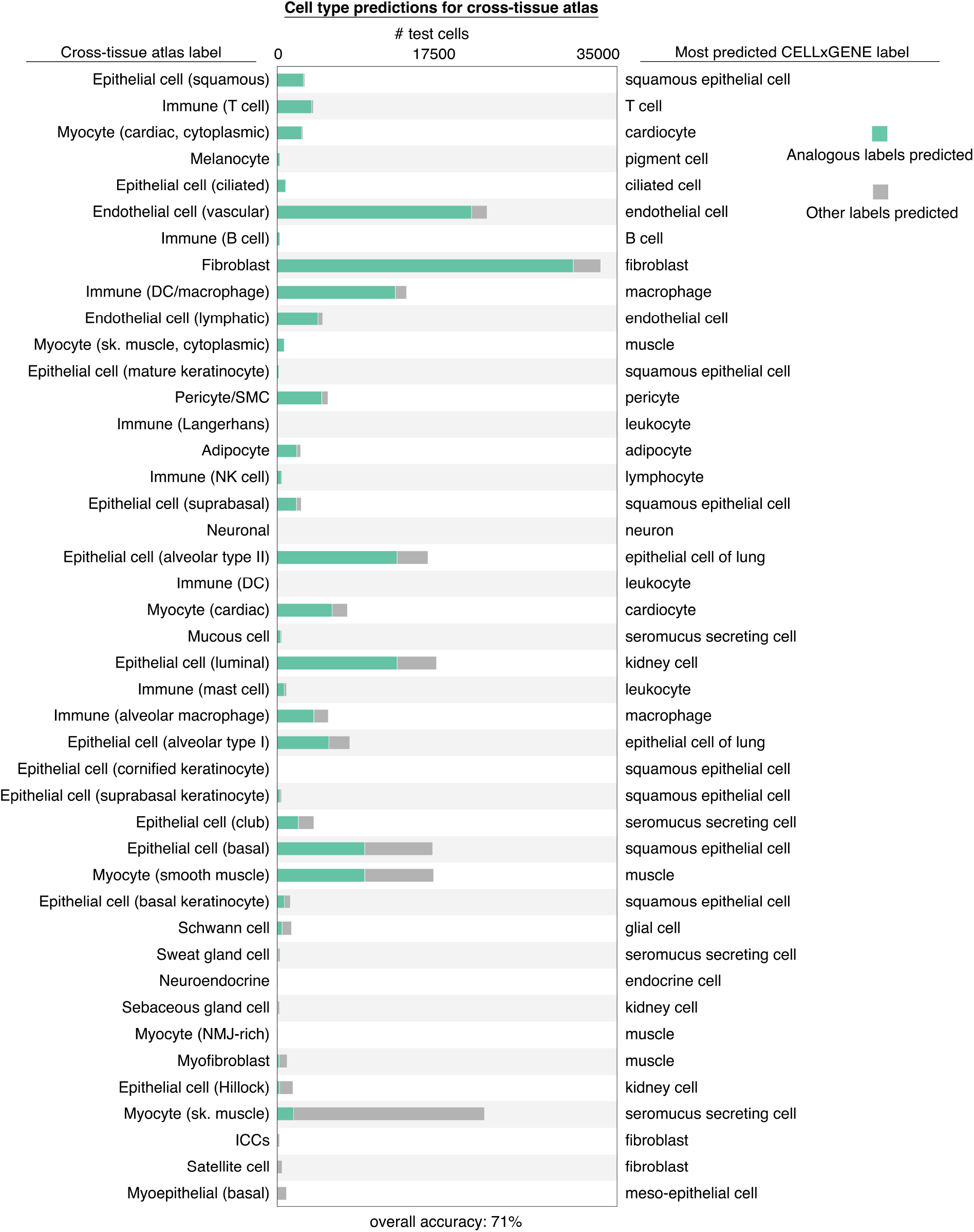
Cell type label transfer to external dataset. Cell type predictions for cells from an external cross-tissue atlas^15^ by the multi-task learning model fine-tuned on CELLxGENE cell types (among other tasks). Cell type labels from authors of the cross-tissue atlas are shown on the left; the most predicted CELLxGENE label for that class is shown on the right. In many cases there was not a direct corollary between cell type labels from the cross-tissue atlas study and the cell type labels used by CELLxGENE. The cell type label translation dictionary used to score correct vs. incorrect label transfer is provided in Extended Data Table 2.

**Extended Data Fig. 4.**
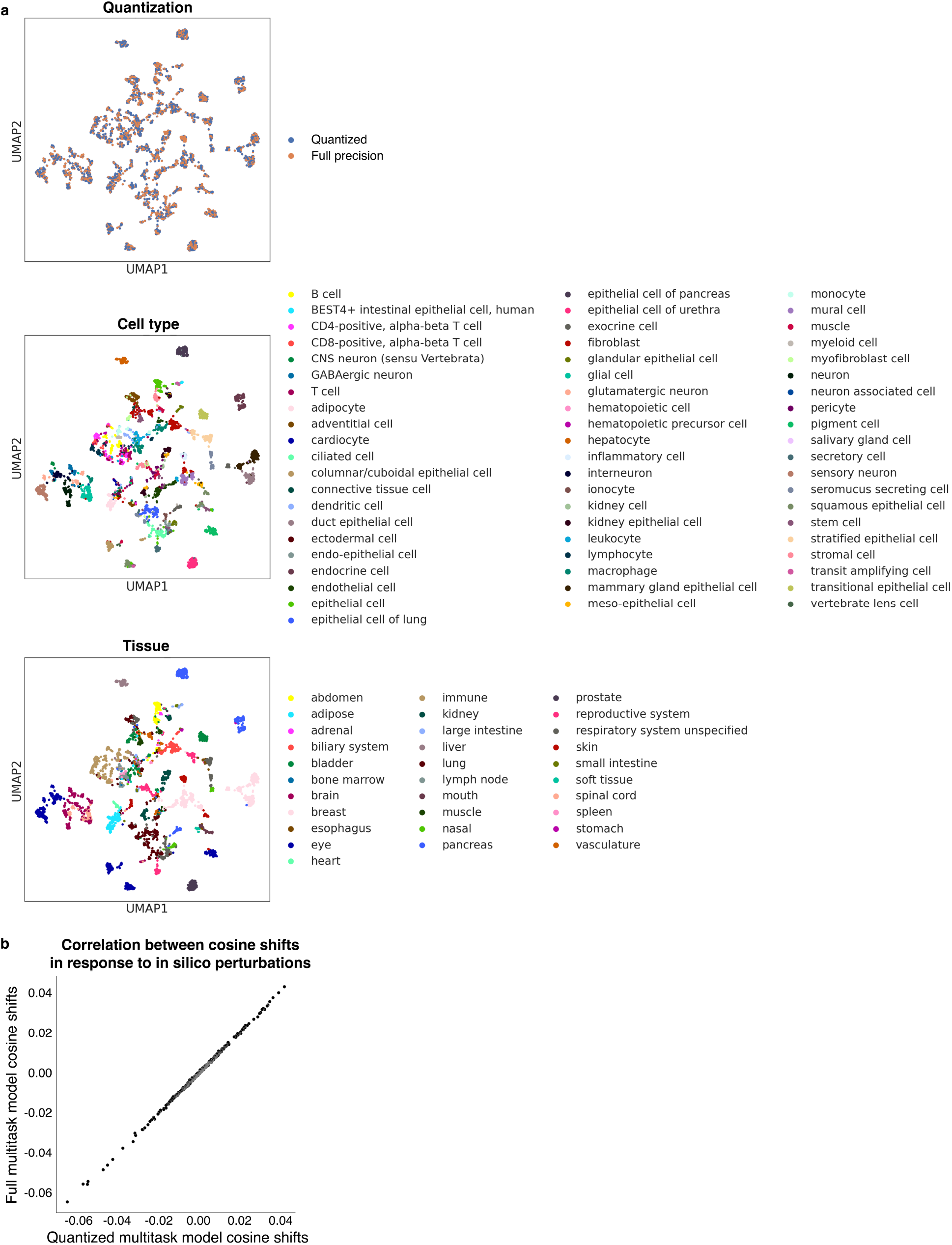
Quantized multi-task model embeddings. **a**, CLS cell embeddings from the 8-bit quantized or full multi-task fine-tuned GF-12L-95M-I4096 of 3000 representative normal adult cells from a broad range of cell types and tissues from the CELLxGENE corpus. **b**, Correlation of in silico treatment analysis results from the 8-bit quantized vs. full multi-task fine-tuned GF-12L-95M-I4096 for an example disease task. Plot shows correlation of cosine shifts from Crohn disease to normal intestinal fibroblasts within the embedding space by in silico deletion of TRRUST genes (each point represents a gene that was in silico deleted in the Crohn fibroblasts) (Pearson correlation 0.9995).

**Extended Data Fig. 5.**
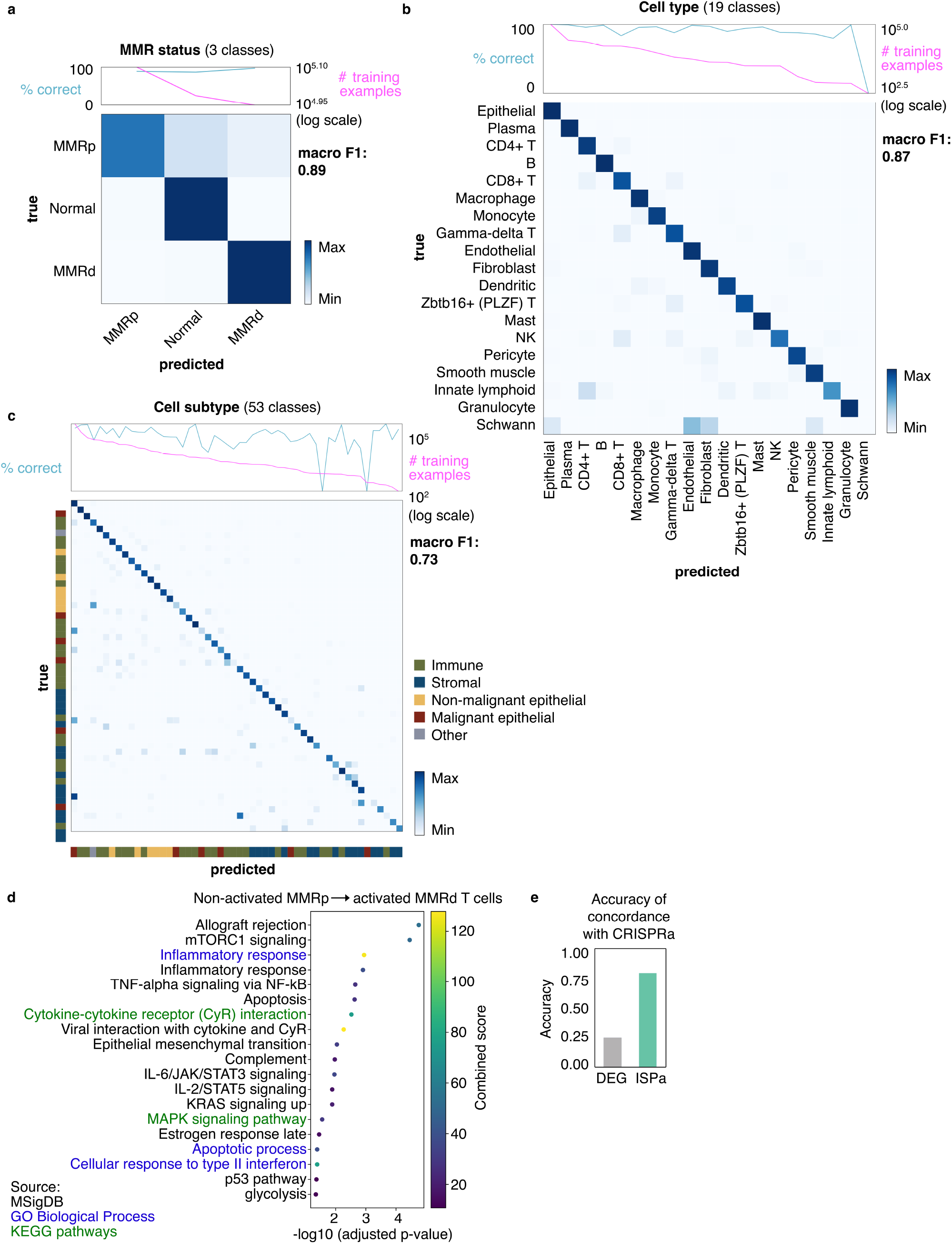
Colorectal cancer multi-task model predictions. **a**, MMR status (3 classes), **b**, cell type (19 classes), and **c**, cell subtype (53 classes) task confusion matrices and macro F1 scores for colorectal cancer multi-task learning model generated by fine-tuning GF-12L-95M-I4096 after cancer domain-specific continual learning with CL max LR rewarm strategy (rewarming (warmup ratio 1%) to the maximum of the learning rate from the initial pretraining and decayed the learning rate on a cosine schedule). Predictions (teal line) were robust to decreasing numbers of training examples (magenta line). Reported results are predictions for individual patients entirely held-out from the training and validation sets. **d**, Gene set enrichment of genes whose in silico activation in non-activated T cells in MMRp tumors were predicted to significantly shift the cell state towards the activated T cell state found in MMRd tumors with higher anti-tumor immune activation. **e**, Accuracy of concordance with CRISPR activation (CRISPRa) screen^24^ for interferon gamma-promoting perturbations in CD8+ T cells. CRISPRa hits were compared to hits from DEG or ISPa analysis in interferon gamma-producing activated T cells from MMRd tumors compared to non-activated T cells from MMRp tumors. Positive hits by ISPa enriched for CRISPRa positive hits by 40% compared to ISPa non-hits. Positively differentially expressed genes were not enriched for CRISPRa positive hits compared to not positively differentially expressed genes. DEG=differentially expressed genes. ISPa=in silico perturbation, activation.

